# Alternate approach to stroke phenotyping identifies a genetic risk locus for small vessel stroke

**DOI:** 10.1101/718221

**Authors:** Joanna von Berg, Sander W. van der Laan, Patrick F. McArdle, Rainer Malik, Steven J. Kittner, Braxton D. Mitchell, Bradford B. Worrall, Jeroen de Ridder, Sara L. Pulit

## Abstract

Stroke causes approximately 1 in every 20 deaths in the United States. Most strokes are ischemic, caused by a blockage of blood flow to the brain. While neurologists agree on the delineation of ischemic stroke (IS) into the three most common subtypes (cardioembolic stroke (CES), large artery stroke (LAS), and small vessel stroke (SVS)), several different subtyping systems exist. The two most commonly-used clinical subtyping systems are TOAST (Trial of Org 10172 in Acute Stroke Treatment) and CCS (Causative Classification System for Stroke), but agreement between these two systems is only moderate. Here, we have compared two approaches to combining the existing subtyping systems for a phenotype suited for a genome-wide association study (GWAS).

We used the NINDS Stroke Genetics Network dataset (SiGN, 13,390 cases and 28,026 controls), which includes cases with both CCS and TOAST subtypes. We defined two new phenotypes: 1) the intersect, for which an individual must be assigned the same subtype by CCS and TOAST; and 2) the union, for which an individual must be assigned a subtype by either CCS or TOAST. The union yields the largest sample size while the intersect may yield a phenotype with less potential misclassification.

We performed GWAS for all subtypes, using the original subtyping systems, the intersect, and the union as phenotypes. In each subtype, heritability was higher for the intersect phenotype compared to the union, CCS (alone), and TOAST (alone) phenotypes. We observed stronger effects at known IS variants with the intersect compared to the other phenotype definitions. In GWAS of the intersect, we identify rs10029218 as an associated variant with small vessel stroke. We conclude that in the absence of a golden standard for phenotyping, taking this alternate approach yields more power to detect genetic associations in ischemic stroke.

**Author summary:** Around one in five people will have a stroke at some point in their life. Most strokes (~80%) are ischemic, caused by a blockage of blood supply to the brain. Ischemic stroke risk is partly influenced by lifestyle, and partly by genetics. There are different ischemic stroke subtypes, and genome-wide association studies (GWAS) indicate that the genetic risk for these subtypes is influenced by different genetic factors. Genetic studies of ischemic stroke are therefore typically performed by analyzing each subtype separately. There are several methods to determine someone’s subtype based on clinical features. To find more genetic factors that influence ischemic stroke risk, we aimed to find a group of patients that are phenotypically similar by using information from all subtyping methods. We compared a group of patients assigned the same subtype by all subtyping methods (the intersect) to a group of patients assigned that subtype by at least one subtyping method (the union). Even though the intersect sample size is smaller, we find genetic factors in the intersect GWAS have stronger genetic effects, likely explained by the fact that we are more certain of the subtype in the intersect. Using the intersect, we find new risk-associated genetic factors.

## Introduction

Stroke is one of the primary causes of death worldwide and causes ~1 in every 20 deaths in the United States [1]. Eighty-seven percent of all strokes are ischemic, caused by a blockage of blood flow to the brain [1]. Ischemic stroke (IS) tends to affect those older than 65 years old and has several known risk factors, including type 2 diabetes, hypertension, and smoking. However, the affected population is extremely heterogeneous in terms of age, sex, ancestral background, and socioeconomic status.

Ischemic strokes themselves are also heterogeneous in terms of clinical features and presumed mechanism. The majority of IS are typically grouped into three subtypes: cardioembolic stroke (CES), typically occurring in people with atrial fibrillation; large artery stroke (LAS), caused by eroded or ruptured atherosclerotic plaques in arteries; and small vessel stroke (SVS), caused by a blockage of one of the small vessels in the brain. These subtypes also seem to be genetically distinct: genome-wide association studies (GWAS) in ischemic stroke have identified single-nucleotide polymorphisms (SNPs) that primarily associate with a specific IS subtype [2]. To date, GWAS have identified 20 loci associated with ischemic stroke, of which 9 appear to be specific to an IS subtype [2]. Furthermore, the subtypes also have varying SNP-based heritabilities (estimated at 16%, 12% and 18% for CES, LAS and SVS respectively [3]), indicating that the phenotypic variation captured by genetic factors varies across the subtypes. Improved genetic discovery can help further elucidate the underlying biology of ischemic stroke as well as potentially help identify genetically high-risk patients who could be candidates for earlier clinical interventions.

While neurologists and researchers agree on the delineation of ischemic stroke into these three primary categories (CES, LAS and SVS), several subtyping systems are currently used to assign a subtype to an ischemic stroke patient. The most commonly used approach is a questionnaire based on clinical knowledge that was originally developed for the Trial of Org 10172 in Acute Stroke Treatment (TOAST) [4]. TOAST was designed for implementation in the clinic and has also been used as subtyping system in the majority of stroke GWAS. More recently, researchers have developed a second subtyping system: the Causative Classification System for Stroke (CCS) [5], a decision model based on clinical knowledge that also incorporates imaging data. There are two outputs of CCS: CCS Causative (CCSc), which assigns one subtype to each patient based on the presumed cause of the stroke; and CCS phenotypic (CCSp), which allows for multiple subtype assignments and incorporates the confidence of the assignment. Previous work indicates that TOAST and CCS have moderate, but not high, concordance in assigning subtypes in patients: agreement is lowest in SVS (κ = 0.56) and highest in LAS (κ = 0.71) [6]. Notably, both subtyping systems still place more than one third of all samples into a heterogeneous ‘undetermined’ category. [6]

Determining a patient’s subtype is difficult and prone to misclassification [7], but critical to genetic discovery in ischemic stroke, as demonstrated by the prevalence of subtype-specific association signals. If a group of cases is comprised of phenotypically heterogeneous samples with different underlying genetic risk, power to detect a statistically significant association at a truly associated SNP is reduced (Fig 1). In contrast, a case definition that captures a more phenotypically homogenous group of cases would improve the chances of detecting genetic variants that associate with disease. Therefore, we used the TOAST, CCSc and CCSp subtype assignments to define two new phenotypes per subtype: the intersect, for which an individual must be assigned the same subtype across all three subtyping systems; and the union, for which an individual must be assigned that subtype by at least one of the subtyping systems. Analyzing the union potentially improves power for locus discovery due to its larger sample size, but at the cost of more potential misclassification. In contrast, analyzing the intersect may improve power for genetic discovery by generating a phenotype that is less prone to mis-classification, despite a smaller sample size.

**Fig 1.**
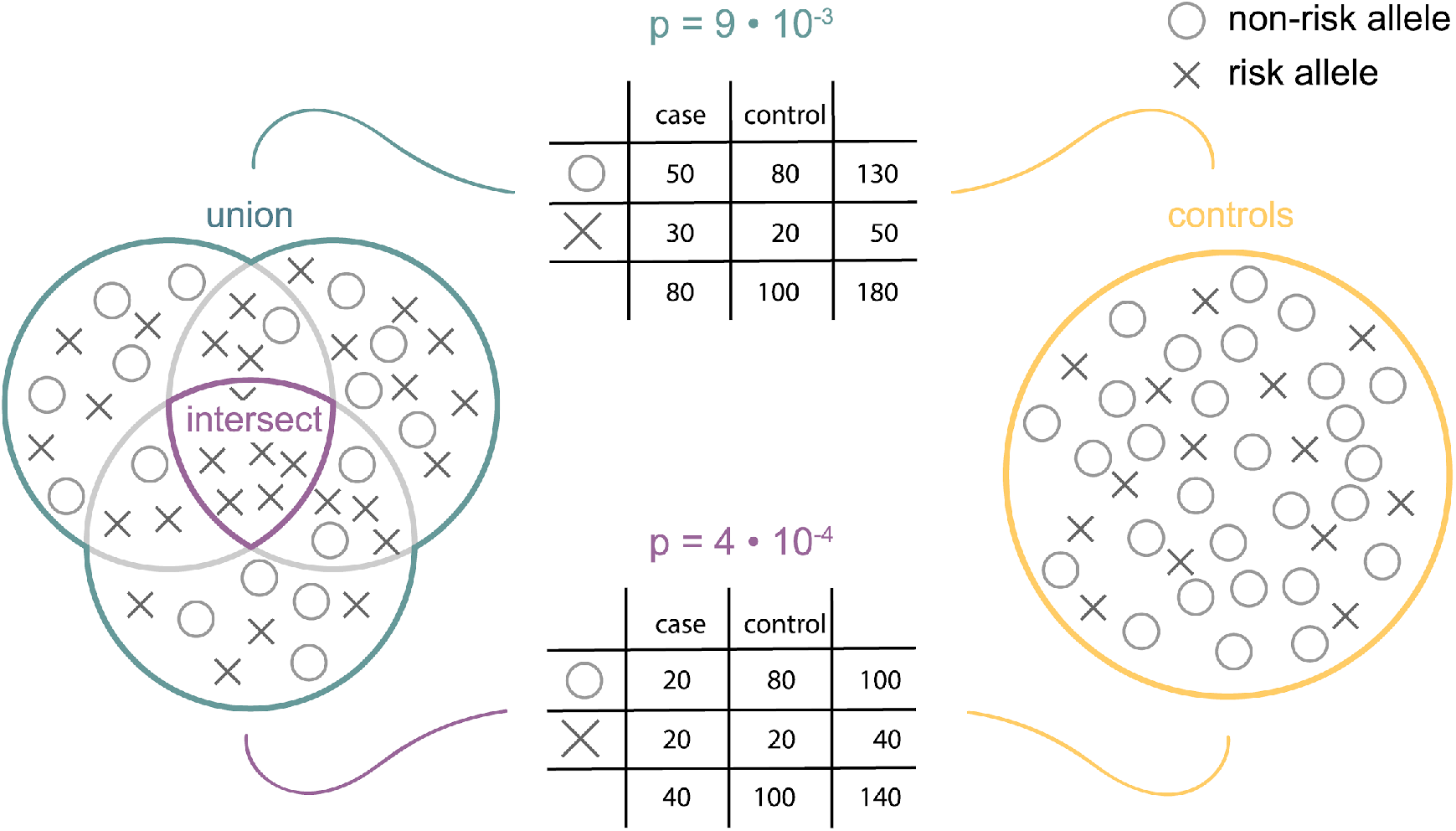
Hypothesized benefit of using the intersect, at a SNP associated with ischemic stroke. Circles indicate the protective allele, and crosses the risk allele. Using a chi-square test (visualized with contingency tables), the measured effect is stronger with a group of cases that is more homogeneous but smaller (intersect, purple) than with a group of cases that is less strictly defined but is larger (union, teal).

Here, we perform GWAS with the union and intersect phenotypes for each primary IS subtype to investigate whether these newly-defined phenotypes indeed improve our ability to detect genetic risk factors for ischemic stroke. We find heritability estimates to be highest in the intersect phenotype for all subtypes. We also find stronger effects at known associations for the intersect compared to the union, and we validate a previously suspected association in small vessel stroke through GWAS of the intersect phenotype.

## Results

### Genome-wide association study data processing

To investigate how redefining stroke phenotypes improves our ability to detect SNPs associated with ischemic stroke, we employed the SiGN dataset. Data processing of the SiGN dataset, including quality control and imputation, has been described in detail elsewhere [8]. Briefly, the dataset includes 13,930 IS cases and 28,026 controls of primarily European descent. Cases and controls were genotyped separately (with the exception of a small number of cohorts) and on various Illumina arrays and then merged together into case-control groups matched for genotyping array and sample ancestry (via principal component analysis). For the cases, phenotype definitions based on one or more of the CCSc, CCSp and TOAST subtyping systems are available (Table 1).

**Table 1.** Case counts for the different phenotype definitions in the three subtypes.

|  | CES | LAS | SVS | undetermined ('other' for CCSp) | total |
| --- | --- | --- | --- | --- | --- |
| **C (CCSc)** | 3,000 | 1,565 | 2,262 | 4,574 | 11,401 |
| **P (CCSp)** | 3,608 | 2,449 | 2,419 | 718 | 9,194 |
| **T (TOAST)** | 3,333 | 2,318 | 2,631 | 3,479 | 11,761 |
| **I (intersect)** | 2,219 | 1,328 | 1,548 | not tested | 5,095 |
| **U (union)** | 4,502 | 3,495 | 3,480 | not tested | 11,477 |
| **S (symmetric difference)** | 2,283 | 2,167 | 1,932 | not tested | 6,382 |

The control group is always the same group of 28,026 individuals

We began our analyses by running genome-wide association studies for all phenotype definitions, including our intersect and union definitions, in all subtypes. We ran all GWAS using a linear mixed model implemented in BOLT-LMM (Supplemental Figure 2) [9]. To take into account any residual population stratification and other batch effects, we included the first 10 principal components and sex as covariates in these analyses (Table S2).

Because the intersect by definition is contained in the union, one additional GWAS for each subtype was run to enable a truly independent comparison of intersect with the symmetric difference (the union minus the intersect). This study focuses on the balance in statistical power between a high sample size and more strictly defined phenotype. Therefore, this sensitivity analysis was only done for the comparison between the two most extreme case definitions.

### Genetic variance in a strictly defined case group explains a higher proportion of phenotypic variance

To estimate how much of the variation in a particular phenotype can be explained by genetic variation, we calculated the heritability (*h*^*2*^) of each phenotype using BOLT-REML, assuming an additive model of effect sizes over all SNPs. We estimated heritability in each of the available phenotypes: the subtypes as defined by TOAST, CCSc, CCSp, the union, and the intersect. We found that the intersect yields a higher *h*^*2*^ estimate than the union in all ischemic stroke subtypes (Fig 2, Table S3). For instance, in CES, h2 of union is 0.139 ± 0.009 and h2 of intersect is 0.275 ± 0.017. We additionally found that the second highest heritability in large artery and small vessel stroke was in CCSc (h2-LAS = 0.258 ± 0.023 and h2-SVS = 0.315 ± 0.029), which assigns only one subtype to each case. The heritabilities for CCSc, CCSp and TOAST were not significantly different from one another in cardioembolic stroke (Table S4), indicating that each original subtyping system is capturing approximately the same proportion of genetic risk.

**Fig 2.**
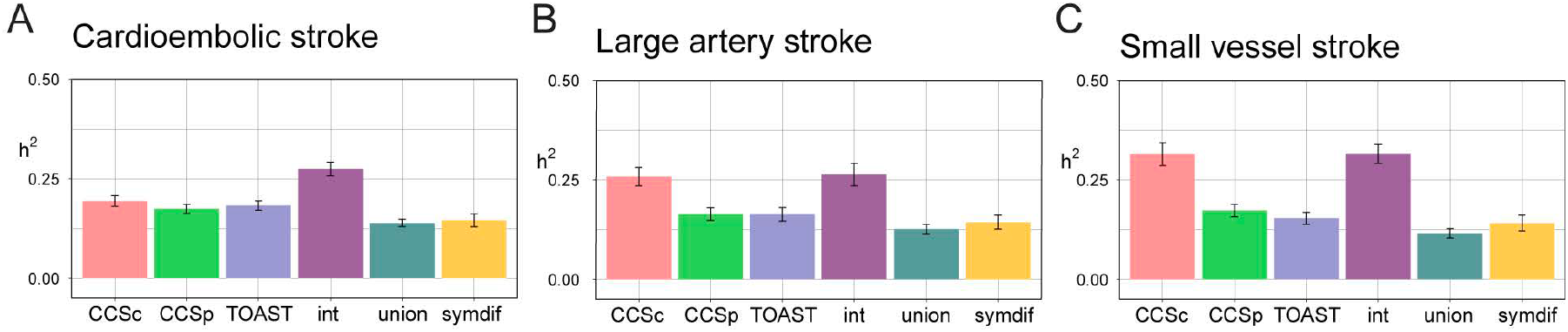
Intersect is the most heritable phenotype. Heritabilities on the liability scale for the six case definitions. int = intersect, symdif = symmetric difference. Bars indicate the 95% confidence interval. (A) In cardioembolic stroke, intersect is significantly more heritable than all other phenotype definitions (p-values for the difference between intersect and all others 3.6e-03 or lower). (B) In large artery stroke and (C) small vessel stroke, intersect is significantly more heritable than all other phenotype definitions except CCSc (p-values for the difference between intersect and all others except CCSc, 2.7e-03 or lower in LAS, 6.1e-07 or lower in SVS). P-values for heritability differences determined by t-test (see Table S5). See Table S4 for numerical values of heritabilities and standard errors.

### Different phenotype definitions represent genetically distinct phenotypes

While heritability gives an estimation of how much variation in a phenotype can be attributed to genetic factors, it does not show how different two phenotypes are from one another (i.e., two phenotypes can have the same heritability and yet be genetically distinct from each other). We therefore evaluated the overlap in significant SNPs for all pairwise combinations of phenotypes for which we performed a GWAS, where high proportions of shared SNPs between two phenotypes indicate genetic similarity. At multiple significance cutoffs, we assessed overlap of significant SNP sets using two complementary similarity measures: the Jaccard index, which measures the ratio of overlapping SNPs (those are significant in both analyses) in the total set of SNPs that are significant in either analysis; and the Pearson correlation of the z-scores of the overlapping SNPs in both analyses (Fig 3). Significance is defined here as an absolute z-score that is higher than the selected z-score threshold (where SNPs can have an effect size < −Z or > +Z). A high Jaccard index indicates that two phenotypes share many of their associated SNPs, while a low Jaccard index means that the phenotypes have distinct genetic architecture. Correlation pertains only to the shared SNPs and indicates if they have similar directionality and magnitude of effect in both analyses.

**Fig 3.**
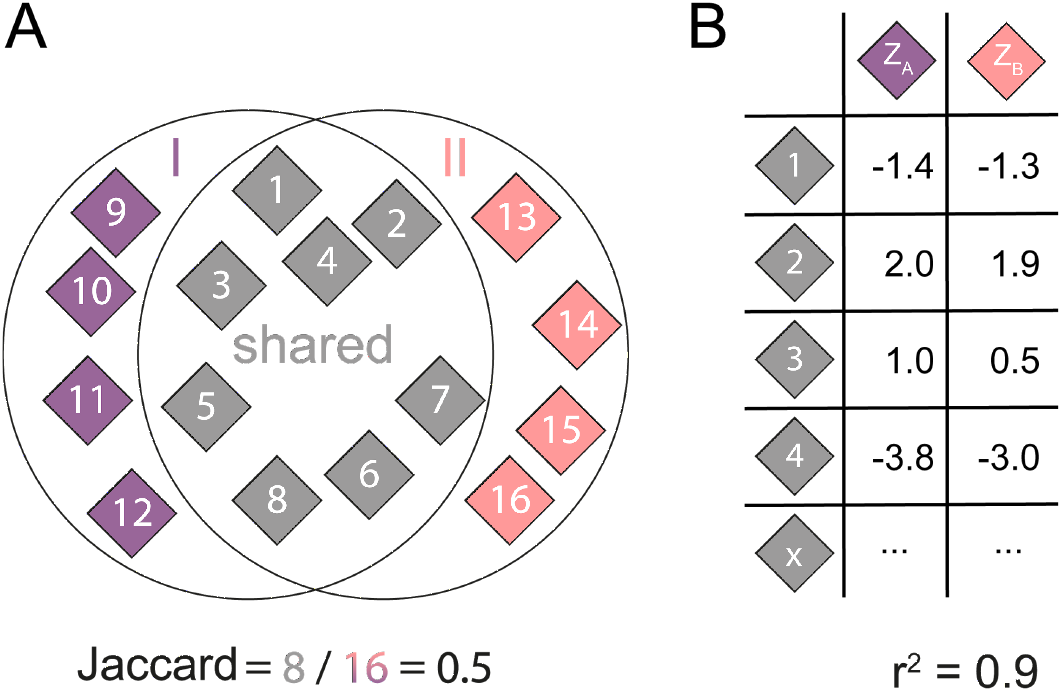
Graphical explanation of overlap analysis. (A) At a certain absolute z-score threshold *Z*, all SNPs that have a z-score lower than −*Z* or higher than +*Z* in analysis I are determined (SNPs 1-8 and 9-12). Next, all SNPs that have a z-score lower than −*Z* or higher than +*Z* in analysis II are determined (SNPs 1-8 and 13-16). The number of shared significant SNPs is divided by the union of significant SNPs to calculate the Jaccard index. (B) We also calculate the Pearson correlation of the z-scores of the shared SNPs.

In order to assess the results of the overlap analyses and their meaning with respect to the ischemic stroke phenotypes, we also performed these analyses between the phenotype definitions and an unrelated GWAS of educational attainment to obtain a null reference (Fig S3).

In cardioembolic stroke (Fig 4, first panel), the Jaccard index for all combinations with intersect decreases with more extreme z-scores to J≈0.2-0.3 while the correlation increases quickly to approach r^2^=1 at Z≈2.5, indicating that a relatively small group of SNPs is significant in both analyses with correlating z-scores, that gets increasingly smaller and stronger correlating. These findings indicate that the stricter the significance threshold is, the fewer shared SNPs there are between any two phenotypes, but that those shared SNPs have more concordant effect sizes. In large artery stroke (Fig S4) and small vessel stroke (Fig S5) the trend is similar, albeit with lower Jaccard indices and correlations, suggesting that there is a set of associated SNPs for each subtype that is found by all phenotype definitions. In all subtypes, when compared to symmetric difference, the intersect is the most genetically distinct phenotype. This confirms that if we combine symmetric difference and intersect, as in the union, we increase phenotypic heterogeneity and thereby decrease the likelihood of detecting a genome-wide significant signal.

**Fig 4.**
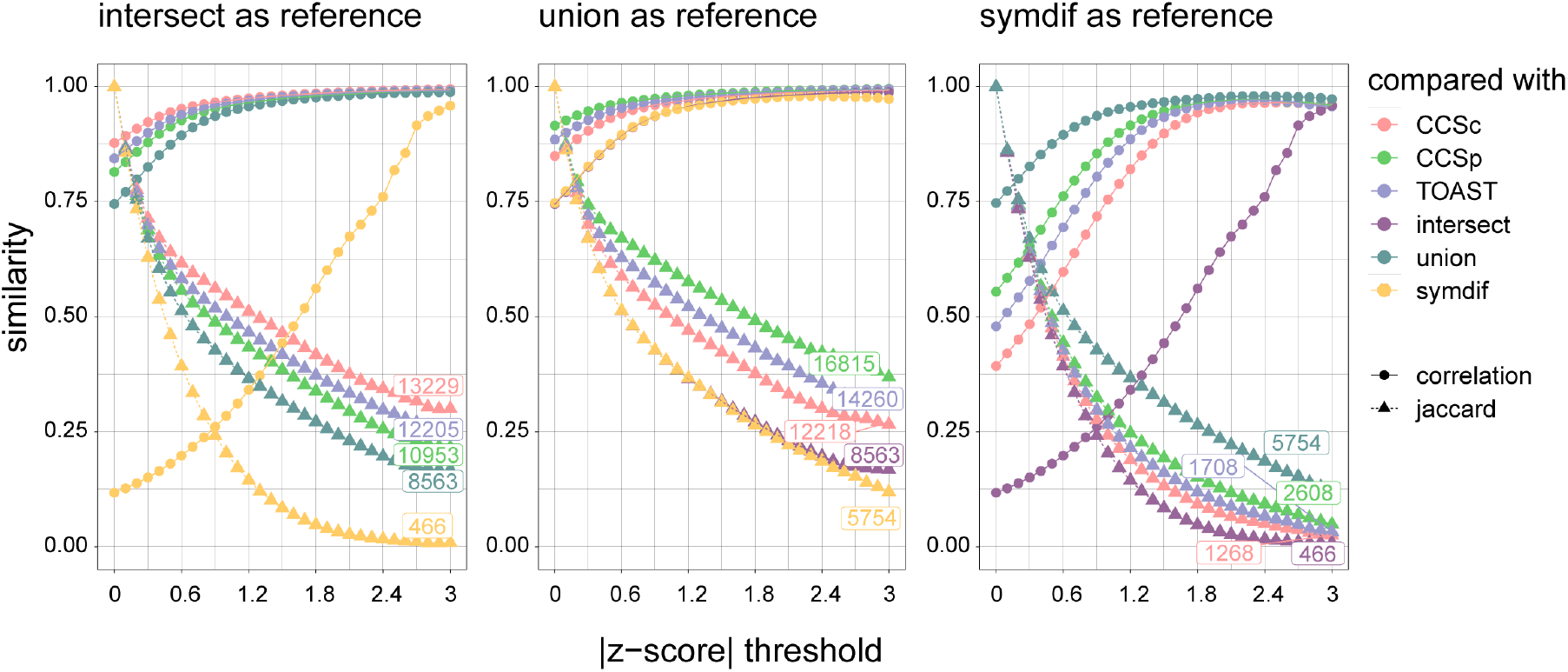
Different phenotype definitions capture different genetic risk factors. Overlap analysis in cardioembolic stroke. Similarity on the y-axis denotes either correlation (circles) or Jaccard index (triangles). The absolute z-score threshold is plotted on the x-axis. Numbers indicate the number of shared SNPs at Z = 3. (A) pairwise comparisons with intersect (B) pairwise comparisons with union (C) pairwise comparisons with symmetric difference.

Fig 4 shows pairwise comparisons only; to investigate if there is one group of SNPs that is significant in all analyses, we also calculated overall Jaccard index: the size of the intersect of SNPs that are significant in all 5 phenotypes (excluding symmetric difference, which we use for sensitivity testing only), divided by the size of their union. The overall Jaccard index (Fig 5) confirms what was suggested by the pairwise overlap analyses: there is a small set of SNPs that is shared across all phenotype definitions, albeit slightly smaller than the pairwise overlapping sets. The Jaccard index is relatively low at higher significance thresholds, indicating that there is also a substantial set of SNPs that is unique to each phenotype definition. Thus, we do find different associated SNPs to ischemic stroke subtypes depending on how exactly the subtype status is defined, but there are some concordant SNPs that are found by all case definitions, regardless of sample size or phenotype homogeneity.

**Fig 5.**
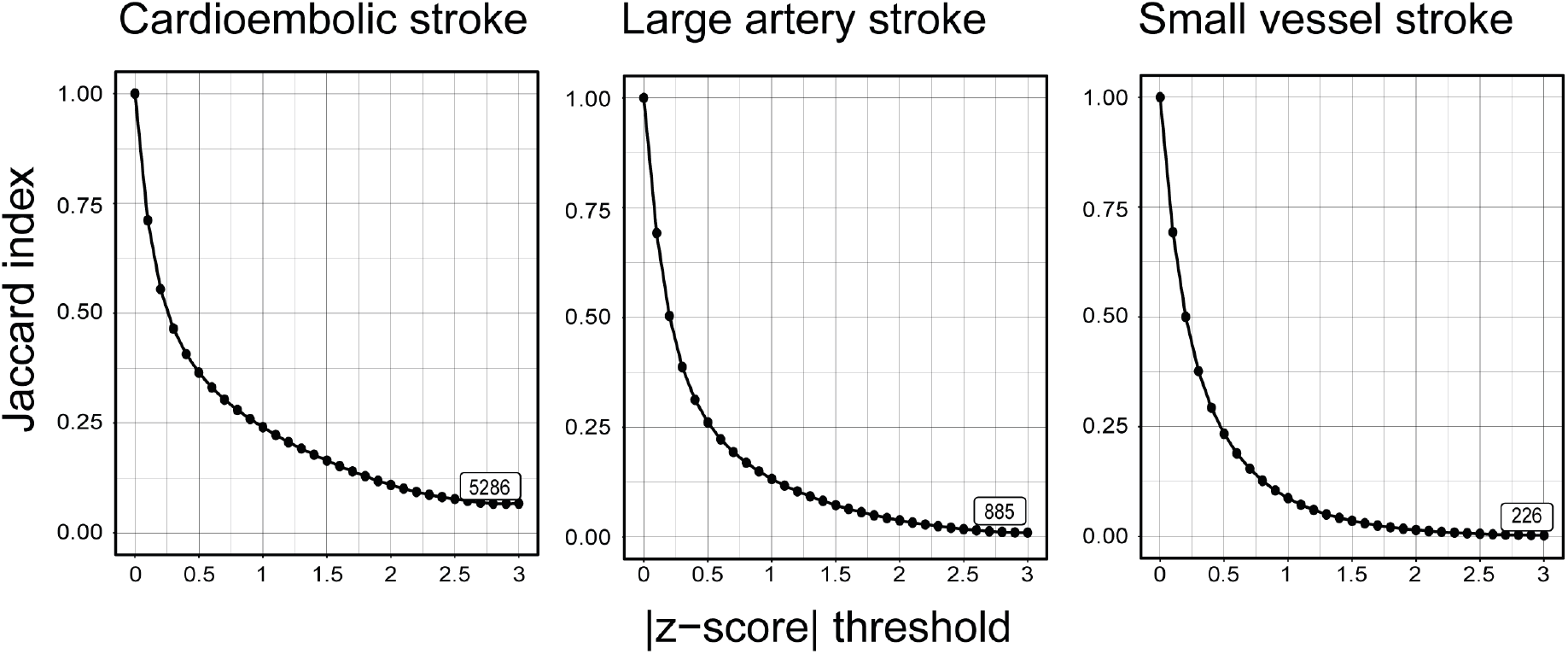
A small set of SNPs is shared between all phenotype definitions. To complement the pairwise overlap analyses, overall Jaccard index was calculated. Jaccard index is plotted on the y-axis, the absolute z-score threshold is plotted on the x-axis. The number of shared SNPs at z = 3 is indicated in the boxes.

### Intersect shows the largest effect at previously known associations

A recent GWAS (MEGASTROKE) in 67,162 TOAST-subtyped cases and 454,450 controls identified 32 loci (22 novel) associated to stroke (either ischemic stroke or intracerebral hemorrhage) and its subtypes [2]. Four of the 32 loci associate to CES, five to LAS, and none to SVS. We investigated the potential to find stroke-associated loci in our redefined phenotypes, with a sample that is 4 to 7 times smaller than MEGASTROKE. To this end, we compared the odds ratios for the 9 known subtype-specific loci in our five phenotype definitions, see Fig 6. In cardioembolic stroke, the intersect phenotype consistently shows the strongest effect. In large artery stroke, intersect shows the strongest effect as well, except at the *LINC01492* locus.

**Fig 6.**
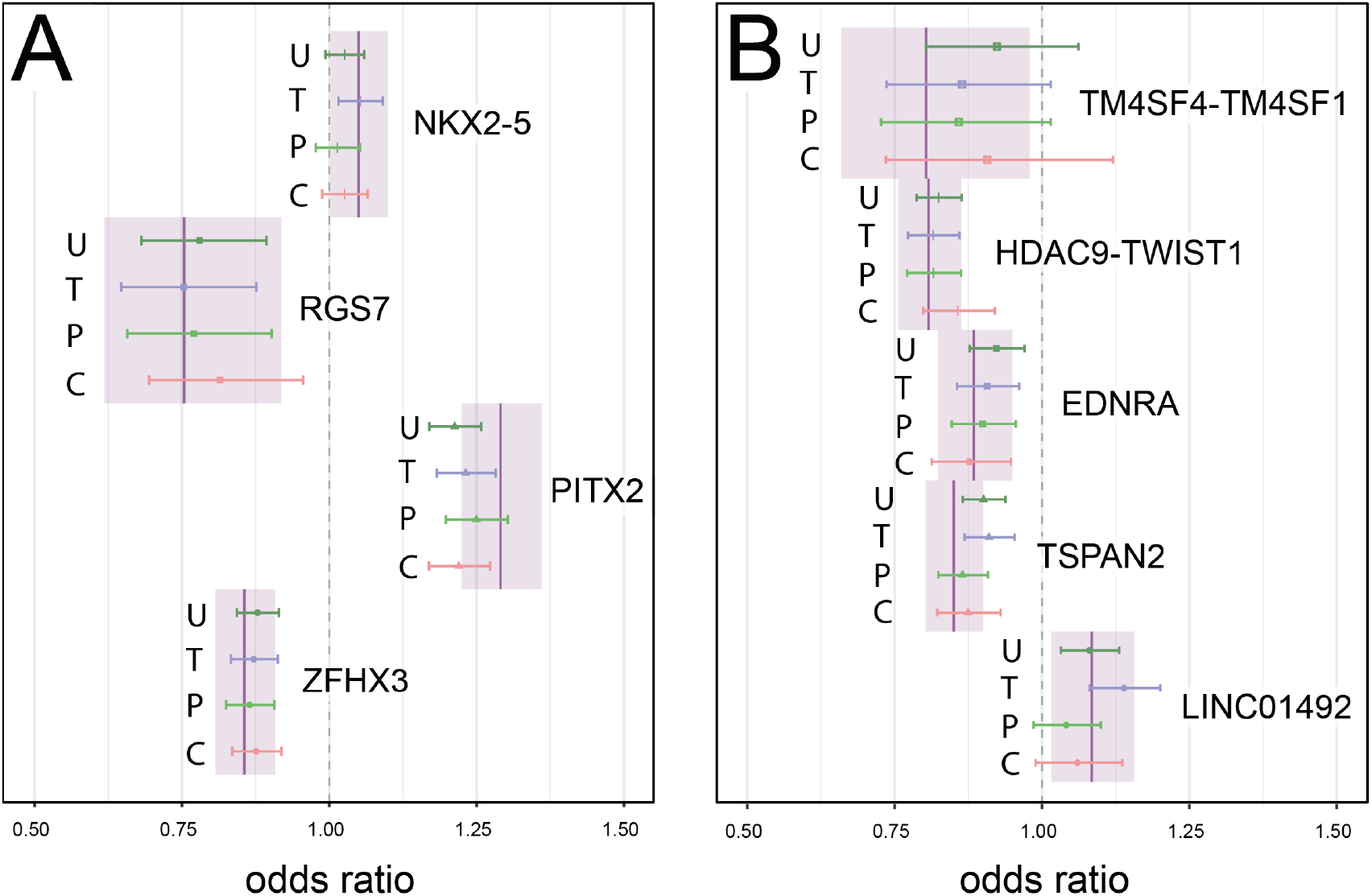
Intersect shows the largest effect at previously identified associations. Odds ratios for (A) the 4 CES-associated SNPs in the five phenotype definitions (B) the 5 LAS-associated SNPs in the five phenotype definitions. U = union, T = TOAST, P = CCSp, C = CCSc. The OR in intersect, with the 95% confidence interval, is indicated with a purple bar and light-purple box. The dotted line indicates an OR of 1 (no effect). Colored points indicate the OR in the corresponding phenotype definition, with error bars indicating the 95 % confidence interval.

Besides comparing the ORs at subtype-specific signals, we also compared ORs at all stroke-associated loci (including any stroke, any ischemic stroke, cardioembolic stroke and large artery stroke), see Fig S1. We found that intersect shows the strongest odds ratio 30 times out of 96, (binomial p = 0.010), indicating that odds ratios derived from the intersect phenotype are indeed stronger than the ORs in the other phenotypes more often than expected by chance.

### A stricter phenotype definition finds a new associated locus to small vessel stroke

Our analyses revealed 5 new loci (2 for SVS and 3 for CES, Table 2) which we validated using data from MEGASTROKE (based on the summary statistics of MEGASTROKE with the SiGN cohort removed, to ensure independence), while correcting for multiple testing per stroke subtype.

**Table 2.** Summary statistics for the new genome-wide significant SNPs.

| Locus | SNP | Chr | A1 | A2 | Analysis | P-value | Freq1 | OR | Beta | SE |
| --- | --- | --- | --- | --- | --- | --- | --- | --- | --- | --- |
| **CAMK2D** | rs10029218 | 4 | A | G | SVS-intersect | 1.20E-08 | 0.12 | 1.27 | 0.02 | 0.00 |
|  |  |  |  |  | SVS-rep-EUR | 2.46E-02 | 0.11 | 1.11 | 0.10 | 0.05 |
|  |  |  |  |  | SVS-rep-TRANS | **5.98E-03** | 0.13 | 1.12 | 0.11 | 0.04 |
| **SH2B3 - BRAP**<br>**- ALDH2** | rs11065979 | 12 | T | C | SVS-CCSp | 9.40E-09 | 0.42 | 1.13 | 0.01 | 0.00 |
|  |  |  |  |  | SVS-rep-EUR | **7.62E-03** | 0.43 | 1.08 | 0.08 | 0.03 |
|  |  |  |  |  | SVS-rep-TRANS | **9.29E-03** | 0.41 | 1.08 | 0.07 | 0.03 |
| PFH20 | rs11697087 | 20 | A | G | CES-intersect | 3.20E-09 | 0.09 | 1.26 | 0.02 | 0.00 |
|  |  |  |  |  | CES-rep-EUR | 1.55E-02 | 0.09 | 1.10 | 0.10 | 0.04 |
|  |  |  |  |  | CES-rep-TRANS | 4.76E-02 | 0.09 | 1.07 | 0.07 | 0.03 |
| 5:114799266 | rs2169955 | 5 | T | C | CES-CCSc | 3.90E-08 | 0.57 | 0.90 | -0.01 | 0.00 |
|  |  |  |  |  | CES-rep-EUR | 1.48E-02 | 0.56 | 0.95 | -0.05 | 0.02 |
|  |  |  |  |  | CES-rep-TRANS | 2.22E-02 | 0.56 | 0.96 | -0.04 | 0.02 |
| **GNAO1** | rs3790099 | 16 | C | G | CES-CCSp | 4.90E-08 | 0.85 | 0.87 | -0.02 | 0.00 |
|  |  |  |  |  | CES-rep-EUR | **2.97E-04** | 0.84 | 0.89 | -0.11 | 0.03 |
|  |  |  |  |  | CES-rep-TRANS | 1.10E-02 | 0.77 | 0.94 | -0.07 | 0.03 |

Per locus, the SiGN GWAS is in the first row, in the format ‘subtype-phenotype’. In the other rows, results in MEGASTROKE are shown with ‘subtype-rep-EUR’ for the Europeans-only analysis, and with ‘subtype-rep-TRANS’ for the trans-ancestry analysis. A1 = Allele 1, A2 = Allele 2, Freq1 = frequency of allele 1, OR = odds ratio, Beta = coefficient, SE = standard error. NB, Beta and SE of SiGN GWAS and MEGASTROKE GWAS are not comparable since they come from linear and logistic regression, respectively. The OR are comparable. We did a Bonferroni correction: for SVS, $\alpha = 0.0125$ and for CES, $\alpha = 0.00625$ . Replication p-values below the threshold are indicated in bold. Two SNPs (rs2169955 and rs146508991, in CES-CCSc) that are relatively close (260 kb) on chromosome 5 were in two different clumps, even though they are in LD ( $r^2 = 0.52$ , $D' = 0.87$ , in a CEU population [10] ) because the distance is just above the threshold (250 kb). Because they are in LD, and just a little farther apart than 250 kb, they were considered to be from the same locus and only the strongest association was kept (rs2169955).

For SVS one variant (rs10029218) in the *CAMK2D* locus (Table 2, Figure S5), was found in the intersect analysis, and replicated in the trans-ancestry analysis of MEGASTROKE. The other SVS associated variant (rs11065979) in the *SH2B3*-*BRAP*-*ALDH2* locus was found in the CCSp analysis, and replicated in both the trans-ancestry analysis and the European analysis (Table 2, Figure S5). For the 3 CES loci, only one variant (rs3790099, in the *GNAO1* gene, found in the CCSp analysis) was replicated in the Europeans-only analysis (Table 2, Figure S5). In a meta-analysis of a) the MEGASTROKE GWAS without the SiGN cohort and b) the SiGN GWAS for these three SNPs, we found consistent direction of effect in both studies and a lower p-value (Table S5).

Previously, one other locus was reported to associate solely with SVS (16q24 [11]). Here, by applying an alternate phenotyping approach, we identify 4p12 as a novel SVS locus. In general, despite the low sample size as compared to MEGASTROKE, we find stronger associations in the intersect GWAS, likely due to the clearer separation of cases and controls.

## Discussion

To help uncover genetic associations with ischemic stroke that as yet have gone undetected, we defined new ischemic stroke phenotypes based on three existing subtyping systems (CCSc, CCSp, and TOAST). Specifically, we studied the intersect and union of these subtyping systems, for all ischemic stroke subtypes. The intersect results in a smaller number of available cases but potentially results in less misclassification due to agreement between subtyping systems. The union is potentially more heterogeneous, but results in a larger available group of cases. We find that the largest proportion of phenotypic variance explained by SNPs is in the intersect phenotype. Further, our overlap analyses show that, for each subtype, the phenotype definitions each have a unique set of significantly associated SNPs, but that there is also a small set of SNPs that is shared among all definitions, with concordant direction of effect and similar trend in magnitude of effect. We also show that the cases that are in the union but not in the intersect, are genetically distinct from the intersect-cases, implying that the union is a combination of phenotypically heterogeneous cases. With an effective sample size that is 4 to 7 times as small as in MEGASTROKE, we find stronger associations (i.e., higher ORs and lower p-values) at known loci by using the intersect (compared to the other phenotype definitions studied here). This indicates that the intersect yields more net power to detect associations due to its stricter definition, despite its lower sample size, and is thus better suited as a phenotype in GWAS.

We identify a previously sub-threshold association with a SNP in an intron of the *CAMK2D* locus in small vessel stroke by using the intersect, further demonstrating the utility of this phenotype in GWAS. *CAMK2D* expresses a calcium/calmodulin-dependent protein kinase [12]; out of all tissues tested in GTEx, the two tissues with the highest expression are both in brain [13]. The *CAMK2D* locus was found to also associate with P-wave [14], an electrocardiographic property that is implicated in atrial fibrillation, a trait that is associated with cardioembolic stroke [3]. Given that the association replicates in an independent dataset, and the protein is expressed in brain, further fine-mapping in this region may give more insight into the biological mechanisms that contribute to stroke. Additionally, we find the *SH2B3 - BRAP - ALDH2* locus to be associated with small vessel stroke. rs11065979 is an eQTL of *ALDH2* (aldehyde dehydrogenase 2) [13]. *ALDH2* is involved in ethanol metabolism; it converts one of the products, ethanal, into acetic acid. The allele that is associated with higher expression of this enzyme is associated with lower incidence of small vessel stroke. *ALDH2* is mainly expressed in liver, but it is also expressed in brain. [13] Previous work has shown an association between higher expression of ALDH2 and lower incidence of stroke in rats [15]*. SH2B3* and *BRAP* are minimally expressed in brain, compared to the other tissues [13]. We also show an association between the *GNAO1* locus and cardioembolic stroke. The protein product of this locus constitutes the alpha subunit of the Go heterotrimeric G-protein signal-transducing complex [12]. It is highly expressed in brain, and while its function is not completely clear, defects in the protein are associated with brain abnormalities [16]. Although this alternate approach to phenotyping has resulted in new associations with two ischemic stroke subtypes, the causality of these loci remains uncertain and warrants further study.

Phenotype definition is an oft-encountered issue in complex trait genetics, as diagnosing and subtyping methodologies can vary and even be contentious within disease areas. Further, phenotype labels are often broad definitions for cases that can be highly heterogeneous when their underlying genetic risk is examined. For example, most psychiatric diseases are also complex and phenotypically heterogeneous, lacking clear and robust diagnostic criteria. In an editorial, the Cross-Disorder Phenotype Group of the Psychiatric GWAS Consortium states: “We anticipate that genetic findings will not map cleanly onto current diagnostic categories and that genetic associations may point to more useful and valid nosological entities”. Our findings here further support this statement, showing that while the original subtyping systems might be useful for diagnosing individual patients, alternative phenotyping approaches and criteria are needed for future genetic studies aimed at unraveling the underlying biology of disease.

## Methods

### The SiGN dataset

The Stroke Genetics Network (SiGN) Consortium composed a dataset consisting of 14,549 ischemic stroke cases. [17] The control group consists primarily of publicly available controls drawn from three large multi-ancestry cohorts. Descriptions of the contributing case and control cohorts have been published previously. [18] Cases and controls have been genotyped on a variety of Illumina arrays, and nearly all cases (~90%) have been subtyped using both TOAST [4] and CCS [19]. All newly-genotyped cases for the latest GWAS are available on dbGAP (accession number phs000615.v1.p1). A previous genome-wide association study has been done on the separate TOAST and CCS subtypes. [18] In this work, we use the same 28,026 controls from this previous GWAS, as well as the 13,930 ischemic stroke cases of European and African ancestry. A third group of cases and controls, primarily comprised of individuals who identify as Hispanic and residing in the United States, has been excluded due to data sharing restrictions. All data processing has been previously described. [18] All genotyping data was generated using human genome build hg19.

### Genome-wide association studies in BOLT-LMM

We ran all GWAS in BOLT-LMM [9], which implements a linear mixed model (LMM). BOLT-LMM implements a Leave-One-Chromosome-Out (LOCO) scheme, so that the genetic relationship matrix (GRM) is built on all chromosomes except the chromosome of the variant being tested. Linear mixed models have been demonstrated to improve power in GWAS while correcting for structure in the data [20]. In addition to the GRM, we included the first 10 principal components as fixed effects. We used the following approximation to convert the effect estimates from BOLT-LMM (on the observed scale) to effect estimates on the liability scale: log(*OR*) = *β*/(*μ* ∗ (1 − *μ*)) where *μ* is the case fraction. [21] For each subtype, the intersect, union and symmetric difference of the original subtyping systems were used as phenotypes in separate GWAS. The original subtyping systems were also used as a phenotype in three additional GWAS per subtype to serve as a point of reference. All ischemic stroke cases that do not belong to the case definition under study were left out of the analysis. The same group of controls is used in all analyses. Association testing was done on all imputed SNPs with a minimum minor allele frequency of 1%. See Supplementary Table 8 in [22] for simulations of type 1 error inflation of BOLT-LMM in datasets with unbalanced case-control ratios. In the GWAS discussed here, case fractions range from 0.05 to 0.14 which means that at variants with MAF >1%, there is no significant inflation of type 1 error rates. Those SNPs that show a large frequency difference (>15%) across the populations in 1000 Genomes were removed (see the methods in [18] for details on how this list of SNPs was compiled). See Fig S2 for QQ-plots (stratified by imputation quality (INFO-score) and by minor allele frequency) and Manhattan-plots. The genomic inflation factor (lambda) varies between 1.0 and 1.1 for cardioembolic stroke and large artery stroke, and between 1.0 and 1.2 for small vessel stroke. We observed a relatively high inflation factor of 1.2 in only the imputed SNPs with a minor allele frequency lower than 5%. Therefore, summary statistics for these SNPs were removed from downstream analyses.

### Heritability estimation in BOLT-REML

To estimate the heritability of the six phenotype definitions for each subtype, we used BOLT-REML [23]. BOLT-REML calculates heritability from the SNPs included in the GRM, and these SNPs must be genotyped (and not imputation dosages). We therefore based our estimates on only genotyped SNPs. Furthermore, we excluded the MHC on chromosome 6, and chromosomal inversions on chromosomes 8 and 17 using PLINK 1.9 [24]. See Table 3 for more information. We filtered on various quality control measures, by passing the following flags to PLINK: --mind 0.05 --maf 0.10 --geno 0.01 --hwe 0.001. Additionally we pruned SNPs at an LD (r^2^) threshold of 0.2 (--indep-pairwise 100 50 0.2). We used the first 10 principal components and sex (determined by presence of XX or XY chromosomes) as covariates. To convert the heritabilities from the observed scale (as if the binary data, coded as 0-1, were continuous) to the liability scale (converting the heritabilities of the observed binary trait to the heritabilities of the underlying, unobserved, continuous liability of the trait), Dempster et al derived a formula that takes into account the prevalence of the disease in the population [25]. In the case of ascertained case-control traits, where the population prevalence is not equal to the study prevalence, this has to be taken into account as well [26]:

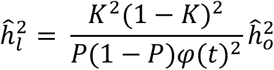

Where 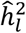 is the heritability on the liability scale, *K* is the population prevalence, *P* is the study prevalence, *t* is the liability threshold, and 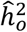 is the heritability on the observed scale. To test for significant difference between the estimated heritabilities, we performed an independent t-test.

**Table 3.** Genomic regions removed before heritability estimation.

| Chromosome | Start (Mb) | End (Mb) | Name |
| --- | --- | --- | --- |
| 6 | 25.8 | 36.0 | MHC |
| 8 | 6.0 | 16.0 | inversion |
| 17 | 40.0 | 45.0 | inversion |

### Overlap analysis

We first calculated z-scores using the following formula: z = beta / se, where beta is the effect size of the SNP and se is the standard error of the beta estimate. The z-scores thus have unit standard error, but we did not standardize them to zero mean (as is the conventional method for calculating z-scores) to maintain the original direction of effect. To assess overlap between two GWAS, we calculated the Jaccard index [27], which is the ratio of a) the number of SNPs significant in both analyses, to b) the number of SNPs significant in either analysis (i.e., the size of the intersect divided by the size of the union of the sets of significant SNPs). The index is a number between 0 and 1: it is 0 if the two sets of significant SNPs do not have any SNPs in common, and it is 1 if the two sets of significant SNPs completely overlap. We additionally calculated, within the set of SNPs that are significant in both analyses, the Pearson’s correlation of the z-scores in the two GWAS to check the concordance of direction and size of effect in the two analyses being compared. Significance was defined as a z-score that is more extreme than an absolute z-score threshold z (varied from 0 until 3, in increments of 0.1). At the most extreme z-score threshold (z>3 or z< −3), the absolute number of SNPs that are significant in both analyses is indicated in the plot. As a null comparator, these overlap analyses were also performed with GWAS results from a study of educational attainment in 1.1 million individuals [28] downloaded from EMBL-EBI’s GWAS catalog. [29] The educational attainment study contains 10,098,325 SNPs, the SiGN study contains 10,156,805 SNPs. The overlap analysis was only done on the SNPs that are present in both datasets: the size of this overlapping set is 7,822,831 SNPs. For the overall comparisons per subtype, we considered all five GWAS. At each z-score threshold, we calculated the overall Jaccard index: the ratio (range between 0 and 1) of the number of SNPs significant in all five analyses to the number of SNPs significant in any analysis. See Fig 3 for a graphical explanation of this method.

### Look-up of Megastroke loci in the union and intersect GWAS

Recently, the MEGASTROKE consortium completed the largest GWAS in ischemic stroke and its subtypes [2]. From this GWAS, we extracted the index SNPs of each genome-wide significant locus in each subtype. We then looked up these SNPs in our GWAS to compare effect sizes, resulting in 15 ORs per SNP (for each of the phenotype definitions in each of the subtypes). See Table S6 for the summary statistics of these look-ups. If the reference allele in MEGASTROKE was not identical to the reference allele in SiGN, the inverse of the odds ratio (1/OR) was taken. We counted how often the intersect showed the most extreme odds ratio, out of all 96 ORs (15 ORs per SNP, for the 32 SNPs that were reported in MEGASTROKE). To determine the probability of the number of times intersect was most extreme, under the null hypothesis that all phenotype definitions are just as likely to show the most extreme OR, we performed a binomial test in R[30].

### Replication of new genome-wide hits in MEGASTROKE

To assess all genome-wide significant loci instead of the individual SNPs, we performed clumping in PLINK 1.9 [24] (http://pngu.mgh.harvard.edu/purcell/plink/). We used all SNPs significant at α = 1×10^−5^ as index SNPs. We generated clumps for all other SNPs closer than 250 kb to the index SNP and in LD with the index SNP (r^2^ > 0.05). We kept clumps if the p-value of the index SNP was lower than 5×10^−8^. From the genome-wide significant clumps, only the unique ones were kept (some clumps significantly associated to multiple case definitions). In the case of duplicates, the summary statistics for the analysis with the lowest p-value were kept. Ambiguous SNPs were removed, and if the reference allele in MEGASTROKE was not identical to the reference allele in SiGN, we calculated the inverse of the odds ratio (1/OR). This resulted in a list of 14 unique SNPs. We checked for SNPs that are not in a locus that had already been reported as an associated locus in MEGASTROKE, resulting in a list of 5 new SNPs (2 for SVS and 3 for CES), which we looked up for replication. To this end, we ran the MEGASTROKE GWAS again (European and trans-ancestry analysis per subtype using TOAST [31]) without the SiGN cohort, to ensure summary statistics independent from the discovery GWAS. We set Bonferroni p-value thresholds to adjust for the number of SNPs looked-up for the phenotype in question, and for the number of GWAS it was looked up in (2, for the European and trans-ancestry analyses). We did a meta-analysis of the MEGASTROKE GWAS without SiGN, and the SiGN GWAS, for the 3 replicating SNPs only (Table S5). We performed meta-analysis in METAL [32], with the inverse-variance weighting scheme.

## Ethics statement

Ethics statement for the original SiGN study can be found in https://doi.org/10.1016/S1474-4422(15)00338-5

