## supplemental tables and figures for "Alternate approach to stroke phenotyping identifies a genetic risk locus for small vessel stroke"

**Supplemental table 1. Sample sizes**

| | | | | | |
|---|---|---|---|---|---|
| LAS | | | | | |
| Phenotype | controls | cases | total | µ (case fraction) | control:case ratio |
| TOAST | 28,026 | 2,318 | 30,344 | 0.08 | 12.09 |
| CCSc | 28,026 | 1,565 | 29,591 | 0.05 | 17.91 |
| CCSp | 28,026 | 2,449 | 30,475 | 0.08 | 11.44 |
| intersect | 28,026 | 1,328 | 29,354 | 0.05 | 21.10 |
| union | 28,026 | 3,495 | 31,521 | 0.11 | 8.02 |
| CES | | | | | |
| Phenotype | controls | cases | total | µ (case fraction) | control:case ratio |
| TOAST | 28,026 | 3,333 | 31359 | 0.11 | 8.41 |
| CCSc | 28,026 | 3,000 | 31026 | 0.10 | 9.34 |
| CCSp | 28,026 | 3,608 | 31634 | 0.11 | 7.77 |
| intersect | 28,026 | 2,219 | 30245 | 0.07 | 12.63 |
| union | 28,026 | 4,502 | 32528 | 0.14 | 6.23 |
| SVS | | | | | |
| Phenotype | controls | cases | total | µ (case fraction) | control:case ratio |
| TOAST | 28026 | 2631 | 30657 | 0.09 | 10.65 |
| CCSc | 28026 | 2262 | 30288 | 0.07 | 12.39 |
| CCSp | 28026 | 2419 | 30445 | 0.08 | 11.59 |
| intersect | 28026 | 1548 | 29574 | 0.05 | 18.10 |
| union | 28026 | 3480 | 31506 | 0.11 | 8.05 |

**Supplemental table 2. Data overview**

| Feature | number of individuals (percentage) |
|---|---|
| **Age (SD)** | 63.93 (16.57) |
| **Chromosomal sex** | 21,152 XX (45%), 25,865 XY (55%) |
| **Hypertension** | 19,835 (54.8%) |
| **Diabetes Mellitus** | 6,846 (18.9% |
| **Atrial fibrillation** | 3,342 (17.5%) |
| **Coronary Artery Disease** | 6,228 (17.7%) |
| **Current smokers** | 6,391 (16.7%) |
| **Former smokers** | 13,203 (34.5%) |

**Supplemental table 3. Heritability estimates from BOLT-REML**

Heritabilities on the liability scale

| Subtype | CCSc | CCSp | TOAST | intersect | union | symdif |
|---|---|---|---|---|---|---|
| CES | 0.194 | 0.174 | 0.183 | 0.275 | 0.139 | 0.146 |
| LAS | 0.258 | 0.164 | 0.164 | 0.263 | 0.126 | 0.144 |
| SVS | 0.315 | 0.173 | 0.154 | 0.316 | 0.116 | 0.142 |

Standard errors of the heritabilities on the liability scale

| Subtype | CCSc | CCSp | TOAST | intersect | union | symdif |
|---|---|---|---|---|---|---|
| CES | 0.013 | 0.011 | 0.012 | 0.017 | 0.009 | 0.016 |
| LAS | 0.023 | 0.016 | 0.017 | 0.029 | 0.012 | 0.017 |
| SVS | 0.029 | 0.016 | 0.015 | 0.024 | 0.012 | 0.020 |

**Supplemental table 4. P-values of heritability differences**

Significant differences at α = 0.005 as determined by t-test (corresponding to a Bonferroni correction for 10 comparisons per subtype), are indicated in the tables below in bold.

| **CES** | CCSc | CCSp | TOAST | intersect | union | symdif |
|---|---|---|---|---|---|---|
| CCSc | 1.0 | 2.6e-01 | 5.3e-01 | **2.2e-04** | **6.5e-04** | 2.2e-02 |
| CCSp | | 1.0 | 6.2e-01 | **1.4e-06** | 1.5e-02 | 1.5e-01 |
| TOAST | | | 1.0 | **1.3e-05** | **3.6e-03** | 6.8e-02 |
| intersect | | | | 1.0 | **5.4e-12** | **7.0e-08** |
| union | | | | | 1.0 | 7.2e-01 |
| symdif | | | | | | 1.0 |

| **LAS** | CCSc | CCSp | TOAST | intersect | union | symdif |
|---|---|---|---|---|---|---|
| CCSc | 1.0 | **8.5e-04** | **1.0e-03** | 8.8e-01 | **3.9e-07** | **8.2e-05** |
| CCSp | | 1.0 | 9.9e-01 | **2.4e-03** | 5.7e-02 | 3.9e-01 |
| TOAST | | | 1.0 | **2.7e-03** | 7.0e-02 | 4.1e-01 |
| intersect | | | | 1.0 | **9.0e-06** | **3.5e-04** |
| union | | | | | 1.0 | 4.1e-01 |
| symdif | | | | | | 1.0 |

| **SVS** | CCSc | CCSp | TOAST | intersect | union | symdif |
|---|---|---|---|---|---|---|
| CCSc | 1.0 | **2.0e-05** | **9.3e-07** | 9.8e-01 | **2.7e-10** | **8.7e-07** |
| CCSp | | 1.0 | 3.7e-01 | **6.1e-07** | 3.7e-03 | 2.1e-01 |
| TOAST | | | 1.0 | **8.5e-09** | 4.7e-02 | 6.2e-01 |
| intersect | | | | 1.0 | **5.4e-14** | **1.7e-08** |
| union | | | | | 1.0 | 2.6e-01 |
| symdif | | | | | | 1.0 |

**Supplemental figure 1. Replication of known loci**

For all previously known hits, odds ratios and their confidence intervals in each of the GWAS described here, are plotted. Colors indicate subtype, symbols indicate phenotype definition.

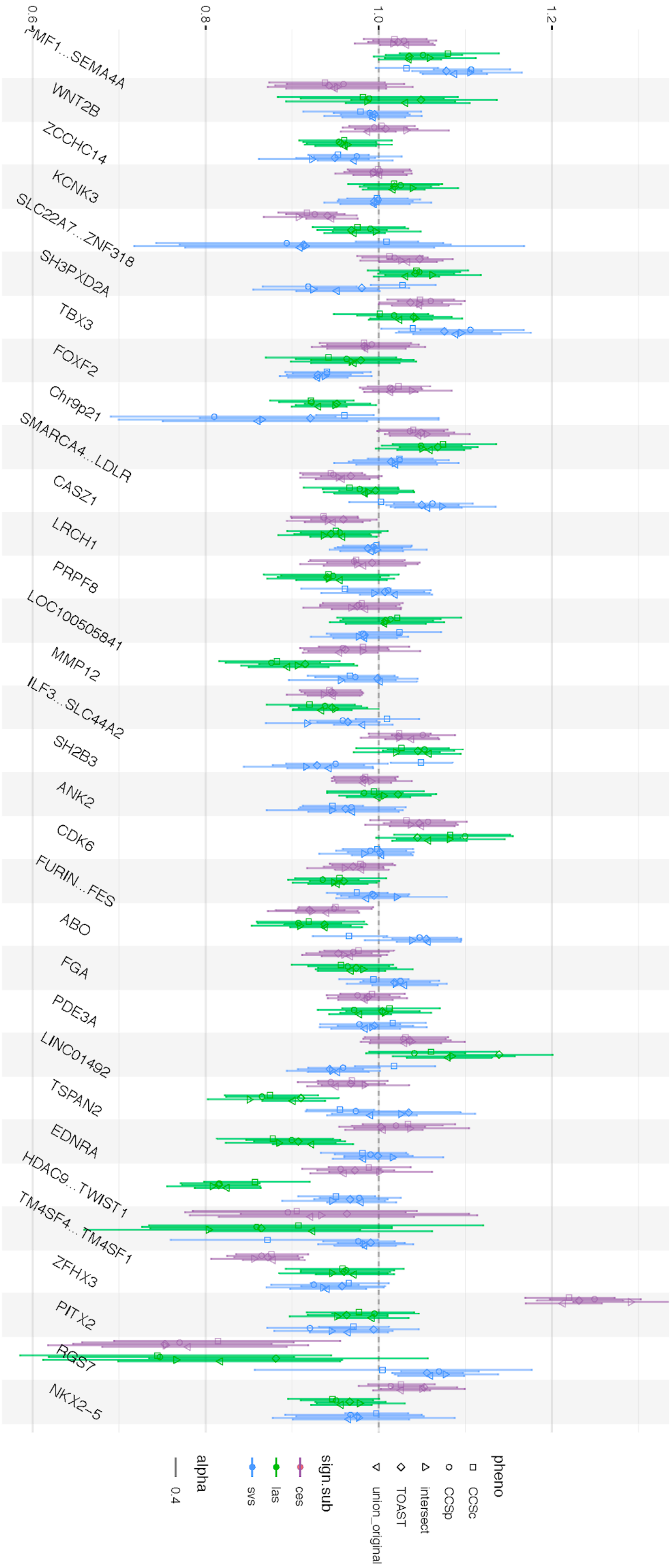

**Supplemental table 5. Meta-analysis of replicated SNPs with MEGASTROKE**

| Chr | Position | MarkerName | A1 | A2 | Freq | FreqSE | MinFreq | MaxFreq | Effect | SE | P-value | Direction |
|---|---|---|---|---|---|---|---|---|---|---|---|---|
| 4 | 114401929 | rs10029218 | a | g | 0.1223 | 0.0003 | 0.1223 | 0.126 | 0.0168 | 0.0029 | 4.30E-09 | ++ |
| 12 | 112059557 | rs11065979 | t | c | 0.4177 | 0.001 | 0.4176 | 0.4302 | 0.013 | 0.0022 | 3.04E-09 | ++ |
| 16 | 56340223 | rs3790099 | c | g | 0.8453 | 0.0002 | 0.8432 | 0.8454 | -0.0206 | 0.0035 | 5.64E-09 | -- |

**Supplemental fig 2. QQ plots and Manhattan plots**

Quality control plots for all GWAS (A) QQ-plot stratified by imputation quality (assessed by INFO score). Inflation factor lambda is indicated in the plot. (B) QQ-plot stratified by minor allele frequency (MAF). Inflation factor lambda is indicated in the plot. (C) Manhattan plot

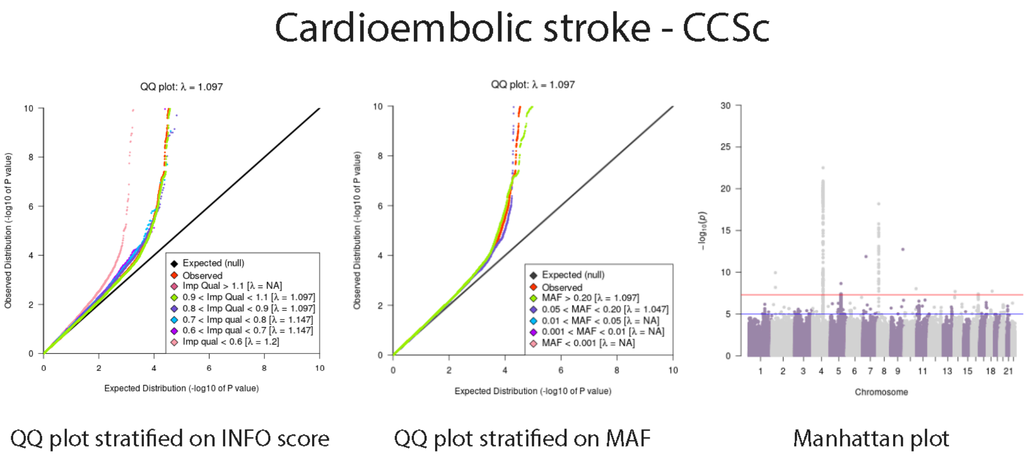

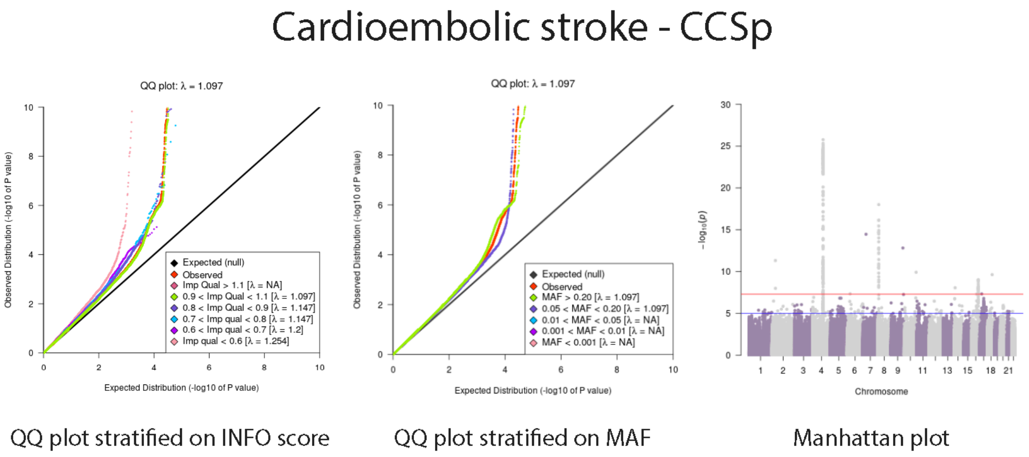

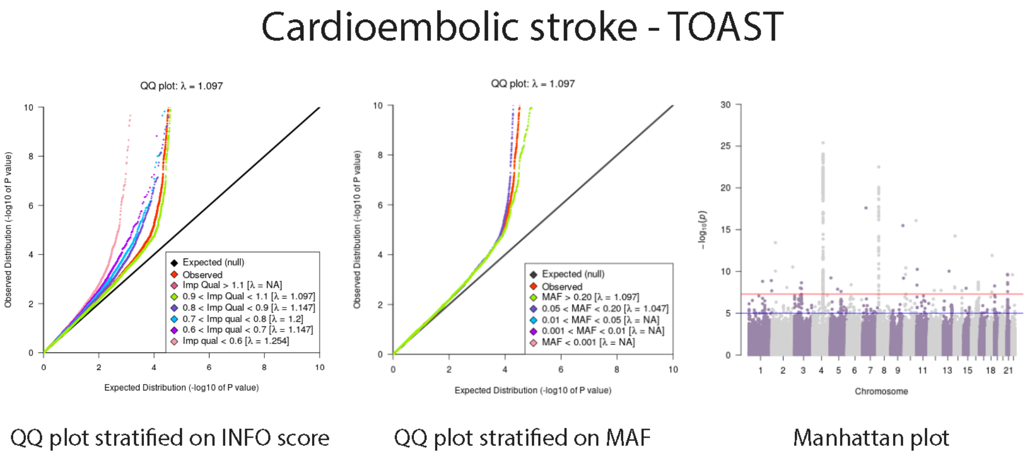

### Cardioembolic stroke - intersect

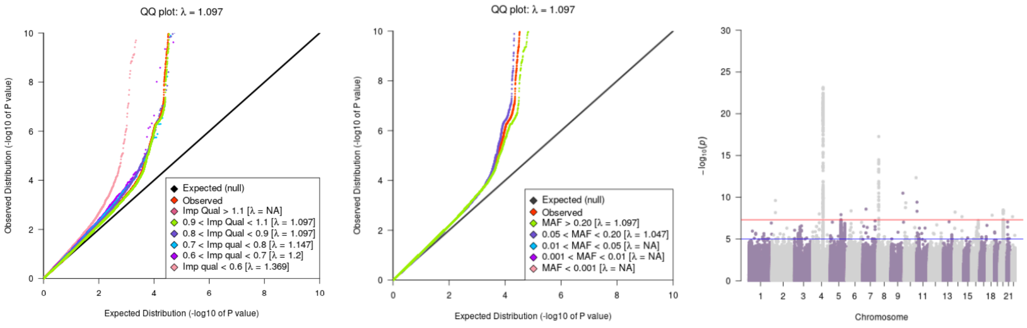

QQ plot stratified on INFO score QQ plot stratified on MAF Manhattan plot

### Cardioembolic stroke - union

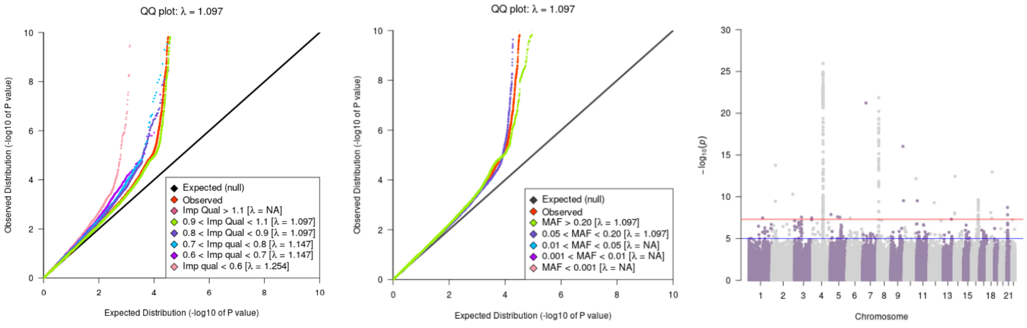

QQ plot stratified on INFO score QQ plot stratified on MAF Manhattan plot

### Cardioembolic stroke - symdif

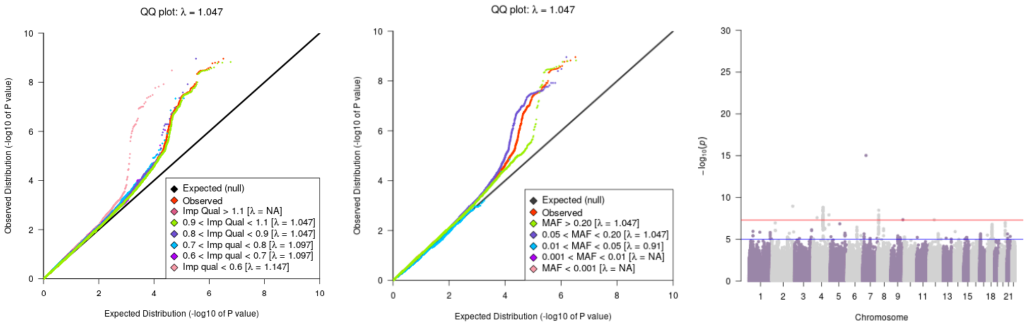

QQ plot stratified on INFO score QQ plot stratified on MAF Manhattan plot

### Large artery stroke - CCSc

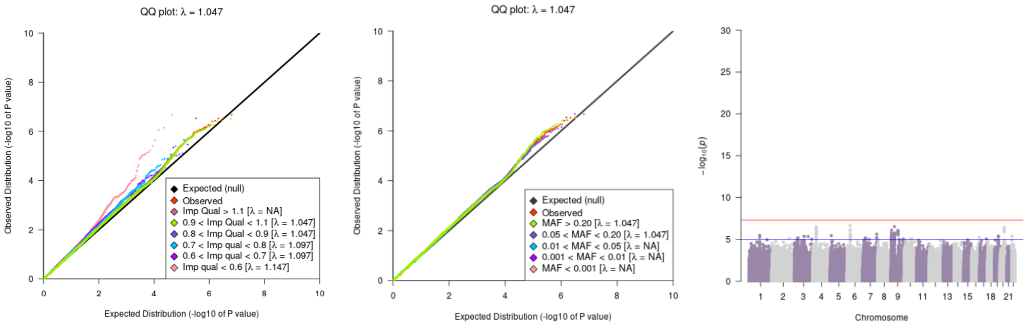

QQ plot stratified on INFO score | QQ plot stratified on MAF | Manhattan plot

### Large artery stroke - CCSp

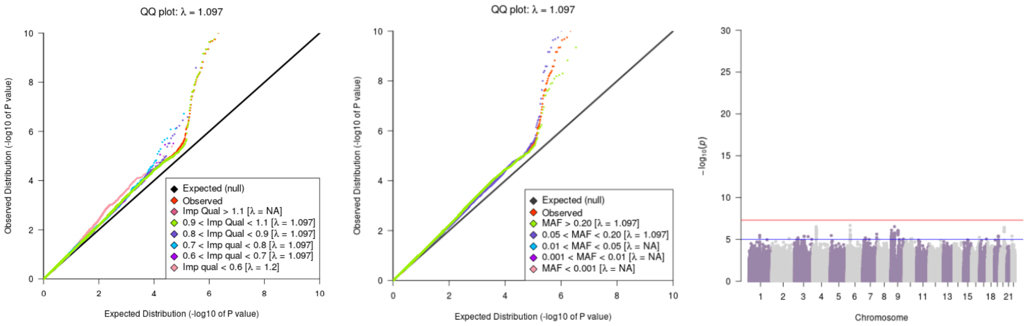

QQ plot stratified on INFO score | QQ plot stratified on MAF | Manhattan plot

### Large artery stroke - TOAST

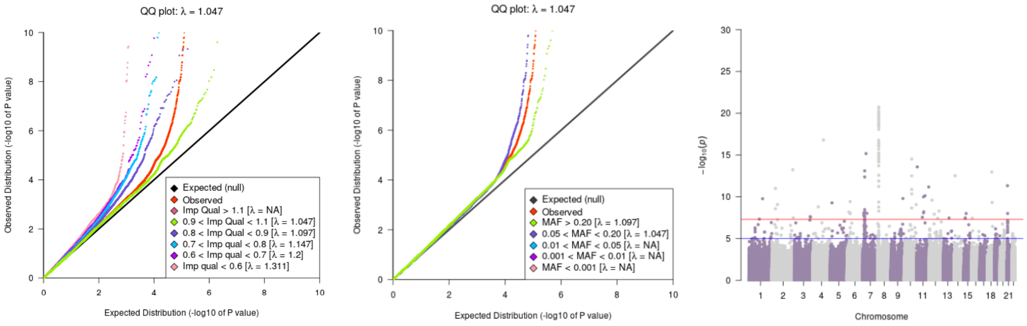

QQ plot stratified on INFO score | QQ plot stratified on MAF | Manhattan plot

### Large artery stroke - intersect

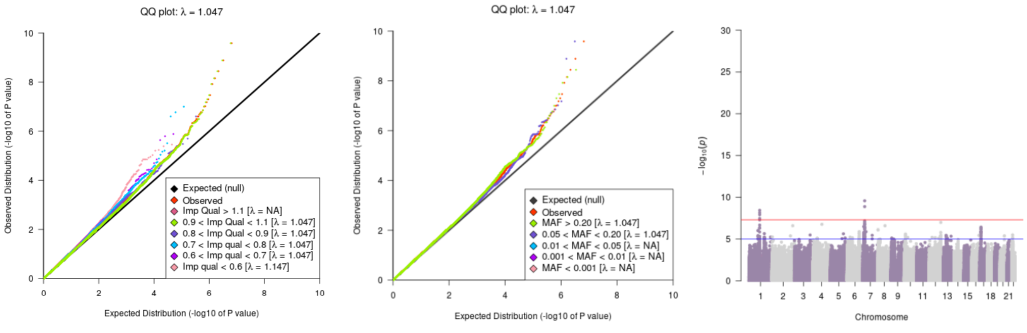

QQ plot stratified on INFO score | QQ plot stratified on MAF | Manhattan plot

### Large artery stroke - union

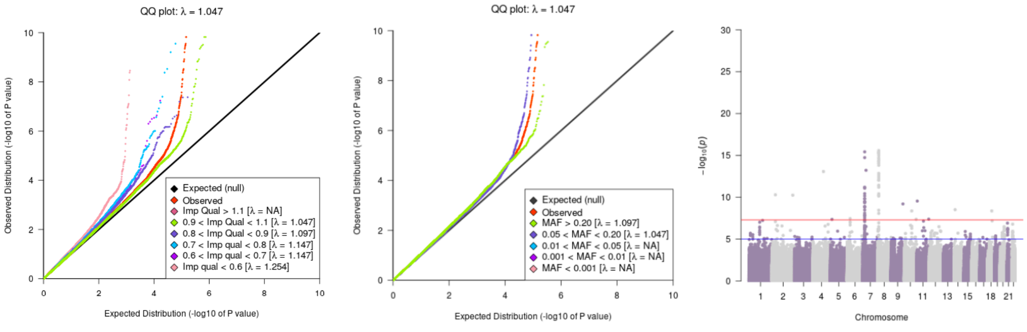

QQ plot stratified on INFO score | QQ plot stratified on MAF | Manhattan plot

### Large Artery Stroke - symdif

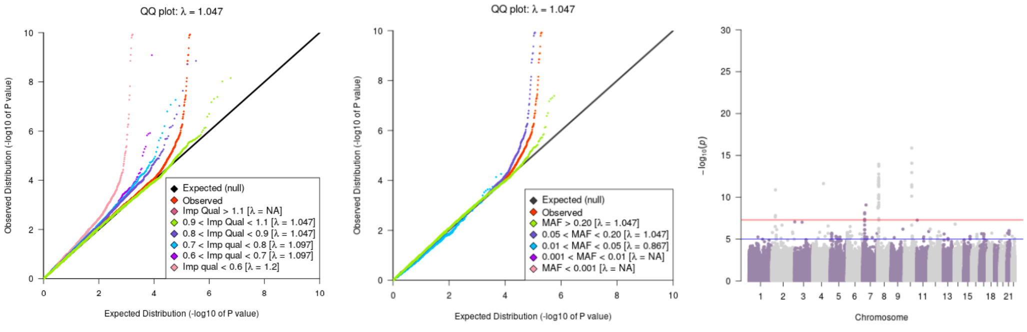

QQ plot stratified on INFO score | QQ plot stratified on MAF | Manhattan plot

### Small vessel stroke - CCSc

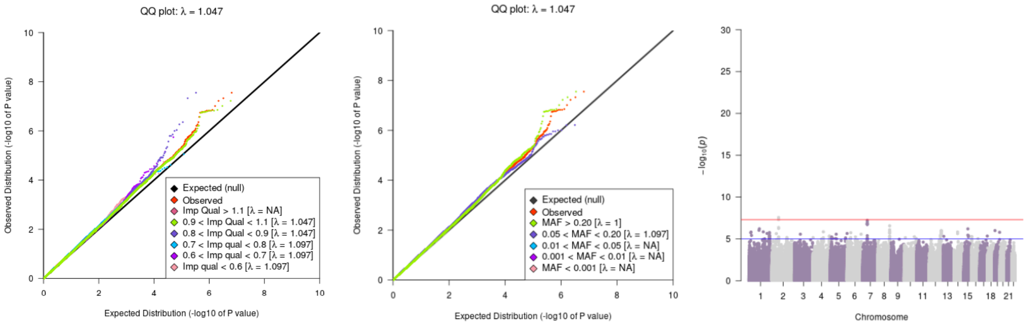

QQ plot stratified on INFO score | QQ plot stratified on MAF | Manhattan plot

50

### Small vessel stroke - CCSp

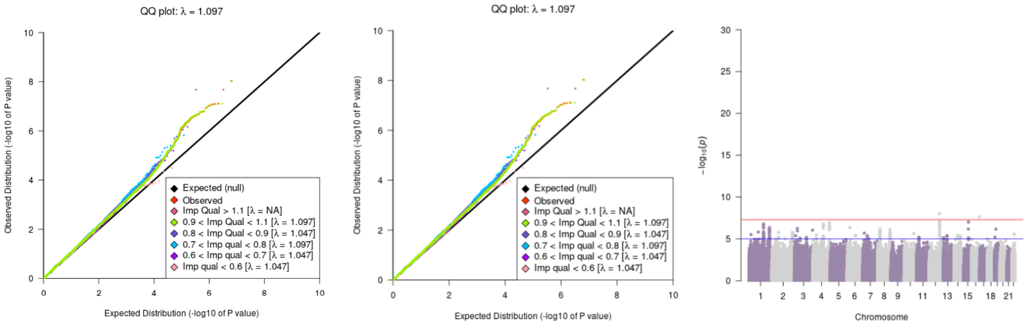

QQ plot stratified on INFO score | QQ plot stratified on MAF | Manhattan plot

51

### Small vessel stroke - TOAST

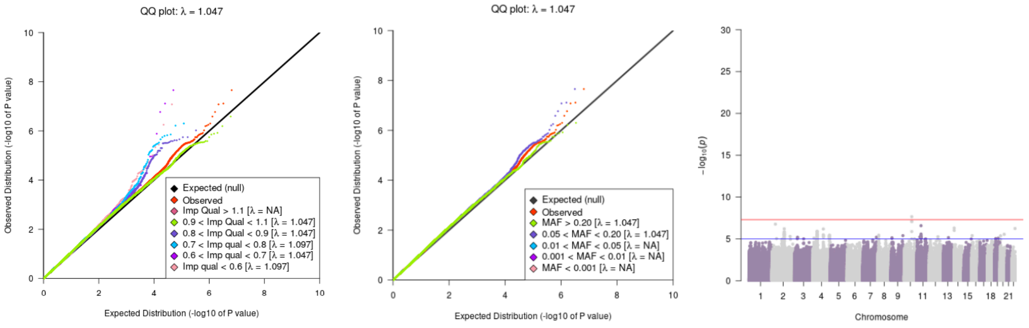

QQ plot stratified on INFO score | QQ plot stratified on MAF | Manhattan plot

52

### Small vessel stroke - intersect

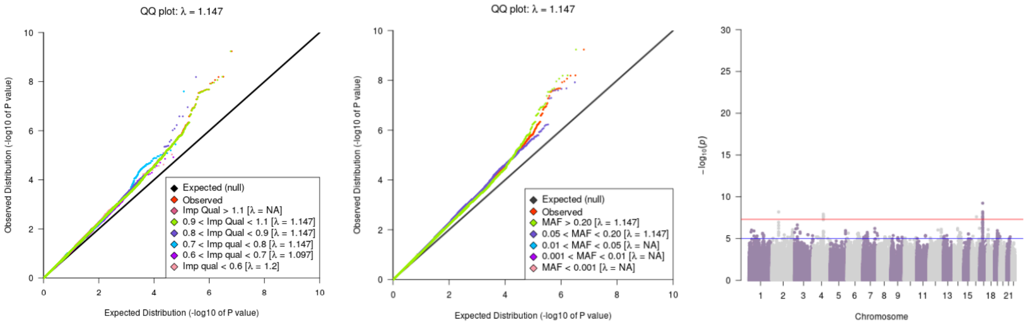

53 QQ plot stratified on INFO score    QQ plot stratified on MAF    Manhattan plot

### Small vessel stroke - union

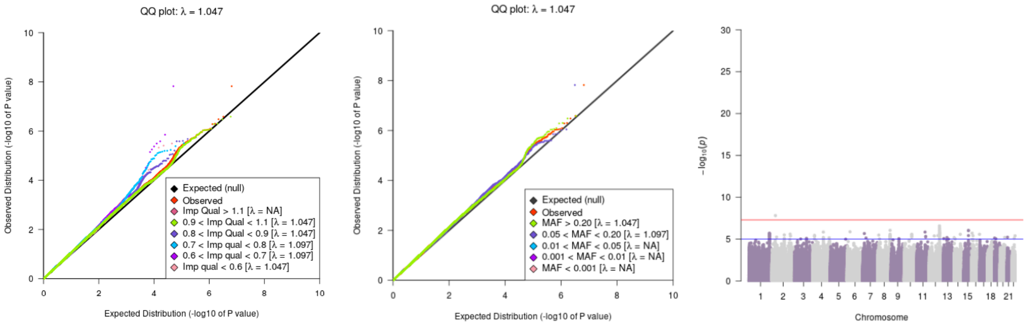

54 QQ plot stratified on INFO score    QQ plot stratified on MAF    Manhattan plot

### Small Vessel Stroke - symdif

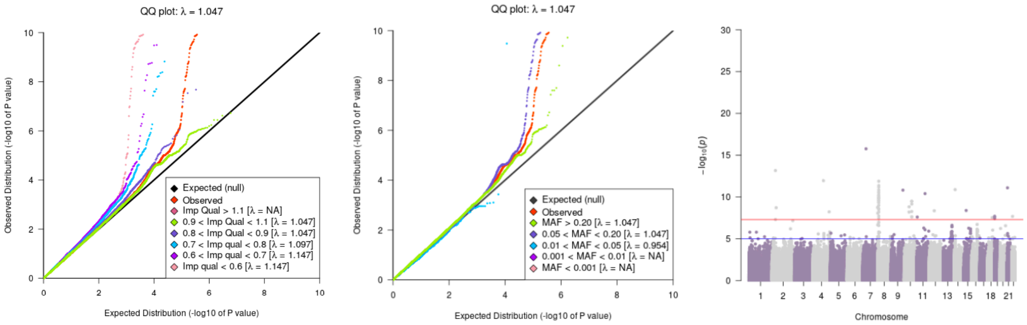

55 QQ plot stratified on INFO score    QQ plot stratified on MAF    Manhattan plot

**Supplemental figure 3. Overlap analysis with cognitive performance as reference**

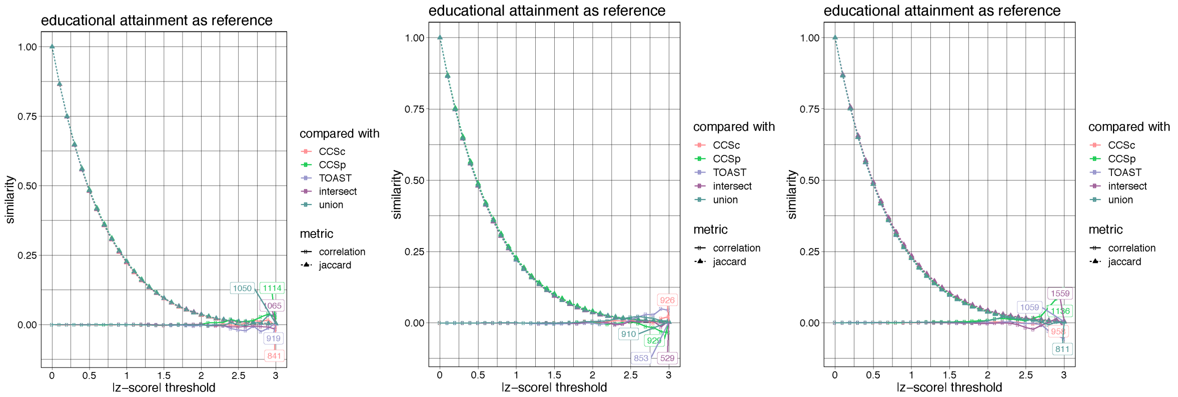

Cardioembolic stroke Large artery stroke Small vessel stroke

**Supplemental figure 4. Overlap plot LAS & SVS**

LAS

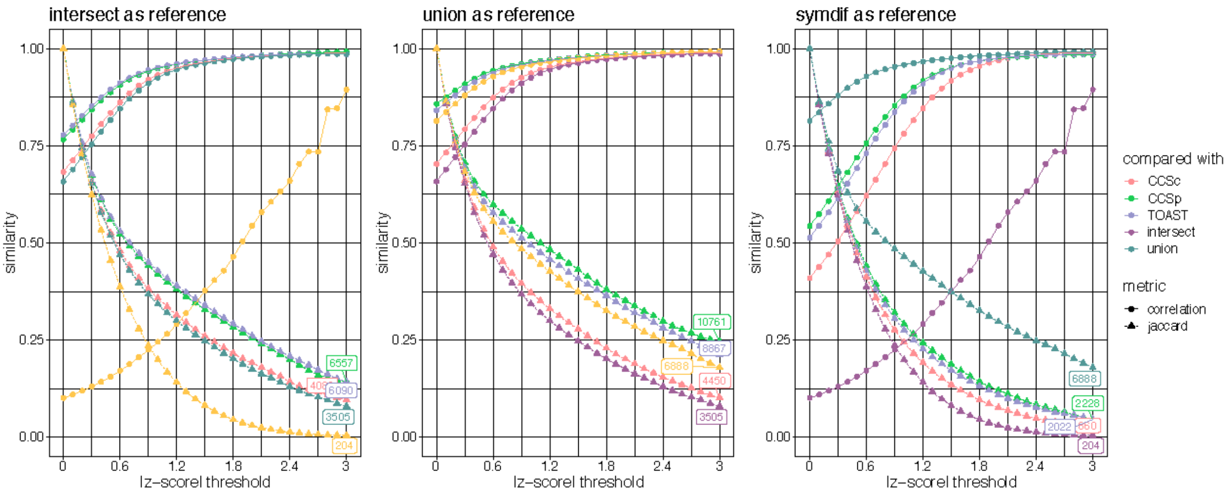

SVS

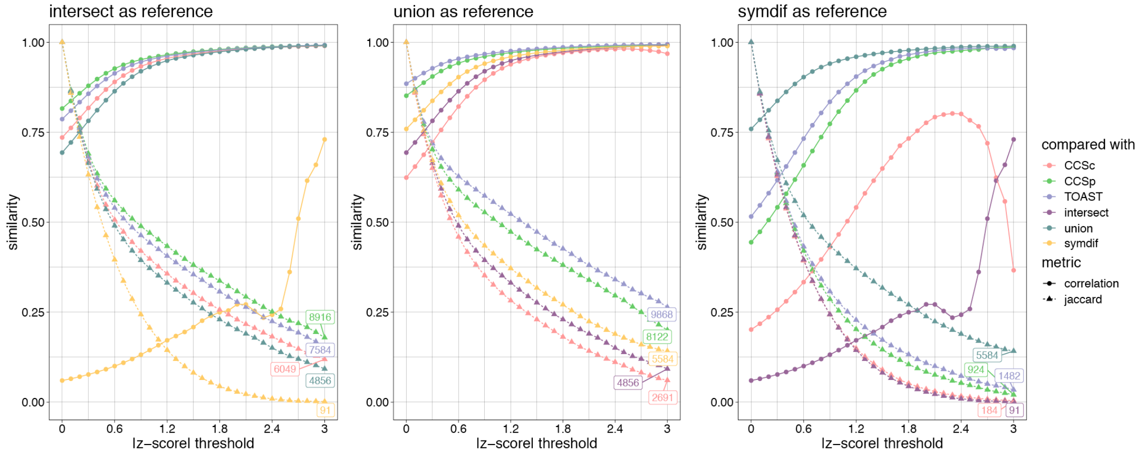

**results of overlap analysis.** Correlation is indicated with points, Jaccard index is indicated with triangles. (A) the other five phenotype definitions compared to intersect (B) the other five phenotype definitions compared to union (C) the other five phenotype definitions compared to symmetric difference

### Supplemental figure 5. Regional association plots

rs11697087

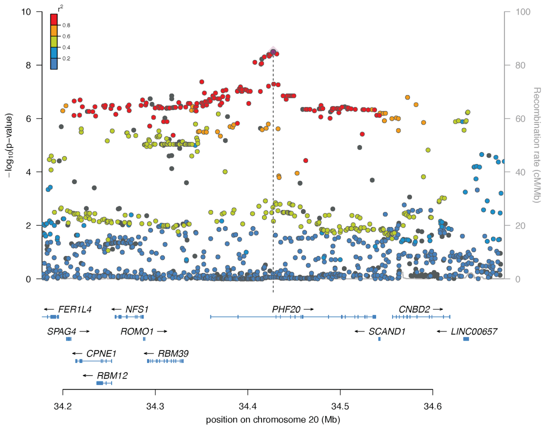

rs11065979

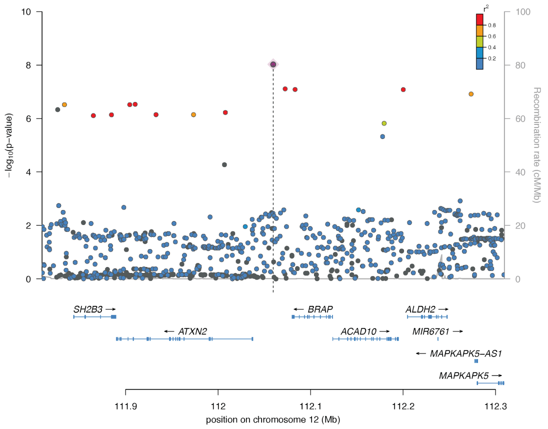

rs10029218

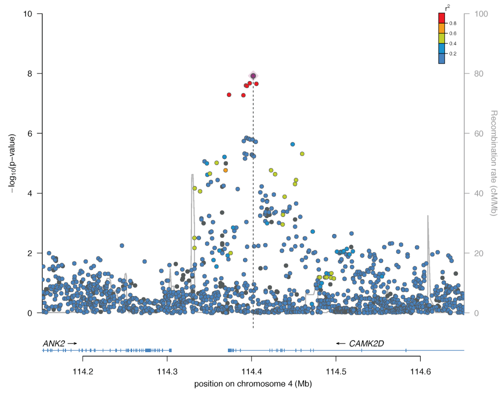

rs3790099

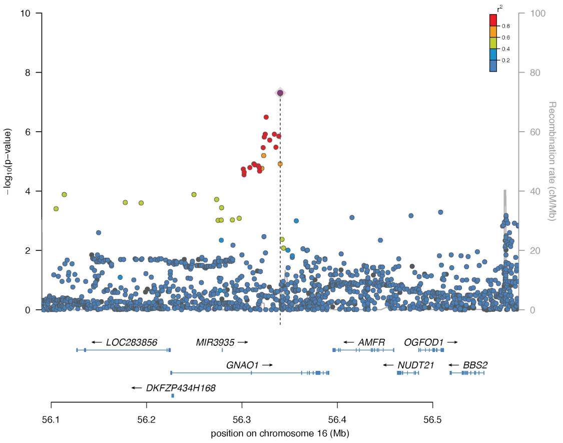

rs2169955

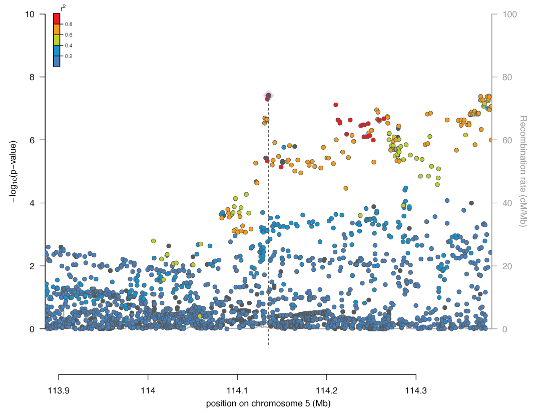

**Supplementary table 6. Summary statistics for previously known associations in the analyses performed in this study**

### See attached files Table-S6.xlsx or Table-S6.ods

**Supplementary Table 7. Summary statistics for previously known associations in the MEGASTROKE study**

### See attached files Table-S7.xlsx or Table-S7.ods
